## Supplementary Material for "AI reveals insights into link between CD33 and cognitive impairment in Alzheimer’s Disease"

### Supplementary Table S1: Module enrichment analysis

**Supplementary Table S1**: **Module enrichment analysis.** If the genes in a module do not enrich NeuroMMSig terms significantly (adjusted $p < 0.05$), individual genes are reported. If significant enriched terms could be found, all significant pathways are reported.

| module | genes | NeuroMMSig term | adj. p-value |
|---|---|---|---|
| 1 | 289 | GABA subgraph | 0.0083 |
| | | Glutamatergic subgraph | 0.0087 |
| 2 | 16 | Toll like receptor subgraph | 0.0089 |
| 3 | 7 | Prostaglandin subgraph | 0.0109 |
| 4 | 6 | TGF-Beta subgraph | 1.4e-05 |
| 5 | 9 | TGF-Beta subgraph | 0.0003 |
| 6 | 4 | Calpastatin-calpain subgraph | 0.0004 |
| 7 | 4 | JAK-STAT signaling subgraph | 0.0393 |
| | | Akt subgraph | 0.0393 |
| | | Cell cycle subgraph | 0.0393 |
| 8 | 3 | AGER, NFATC1, CSF2 | |
| 9 | 3 | Chaperone subgraph | 0.0396 |
| | | Chemokine signaling subgraph | 0.0396 |
| 10 | 2 | REL, IL21 | |
| 11 | 2 | Ubiquitin degradation subgraph | 0.0109 |
| | | Unfolded protein response subgraph | 0.0217 |
| 12 | 2 | Insulin signal transduction | 0.0359 |
| | | Glutamatergic subgraph | 0.0359 |
| 13 | 2 | Peroxisome proliferator activated receptor subgraph | 0.0024 |
| 14 | 2 | GDNF, CASP3 | |
| 15 | 2 | Gamma secretase subgraph | 0.0335 |
| | | Notch signaling subgraph | 0.0335 |
| | | ADAM Metallopeptidase subgraph | 0.0335 |
| 16 | 2 | Epigenetic modification subgraph | 0.0109 |
| 17 | 2 | TICAM1, RALBP1 | |
| 18 | 2 | Amyloidogenic subgraph | 0.0145 |
| 19 | 2 | Tumor necrosis factor subgraph | 0.0430 |
| | | Calcium-dependent signal transduction | 0.0430 |
| | | Unfolded protein response subgraph | 0.0430 |
| 20 | 2 | Acetylcholine signaling subgraph | 0.0205 |
| 21 | 2 | Matrix metalloproteinase subgraph | 0.0302 |
| 22 | 2 | Axonal guidance subgraph | 0.0121 |
| 23 | 2 | IFNB1, TRAF1 | |
| 24 | 2 | XIAP subgraph | 0.0114 |
| | | T cells signaling | 0.0324 |
| 25 | 2 | GPR3, ARRB2 | |
| 26 | 2 | Endoplasmic reticulum-Golgi protein export | 0.0288 |
| | | Amyloidogenic subgraph | 0.0288 |
| 27 | 2 | Low density lipoprotein subgraph | 0.0193 |
| 28 | 2 | MIR485, DLG4 | |
| single genes | 4 | CD33, HSPB2, HSPB3, MIR101-1 | |

#### Supplementary Table S2: Module assignment

The full list of module assignment of each gene in the knowledge graph can be found here and in the attached table. For every gene the corresponding module number from the Markov clustering is given. Module 0 refers to all standalone genes.

#### Supplementary Note S1: iVAMBNs Module Definition

The modules for the VAMBN network are defined through the clustering of the knowledge graph. There were two exceptions: First, modules 4 and 5 enrich the same NeuroMMSig mechanism, namely, TGF-beta signaling, and were thus merged into one module for modeling. The second exception is based on the unavailability of some genes in the gene expression data: Out of the total of 383 genes in the knowledge graph, only 330 were expressed in all AD cohort studies, resulting in seven modules consisting of only one gene after mapping to the gene expression data. Therefore, these genes were used as standalone genes in the VAMBN approach. Altogether there were eight standalone genes: CASP7, CD33, DLG4, GRIN1, NAV3, PPARG, REL and TRAF1. Genes HSPB2, HSPB3 and MIR101-1 were not be measured in all datasets.

On top of modules comprising molecular mechanisms, one *phenotype* module summarizes MMSE scores and Braak stages and some demographic features, namely age, gender, years of education, the APOE genotype, and the brain region are integrated as standalone features.

After module definition, the next step in iVAMBN model generation, is the training of the auto-encoded values per module. Therefore, a HI-VAE was trained for each module separately. A grid search was used to find the best hyperparameters, and each candidate hyperparameter set was evaluated via a 3-fold cross validation. Once finished with this process, each module dimension was reduced to one.

#### Supplementary Figure S1: Clustered knowledge graph.

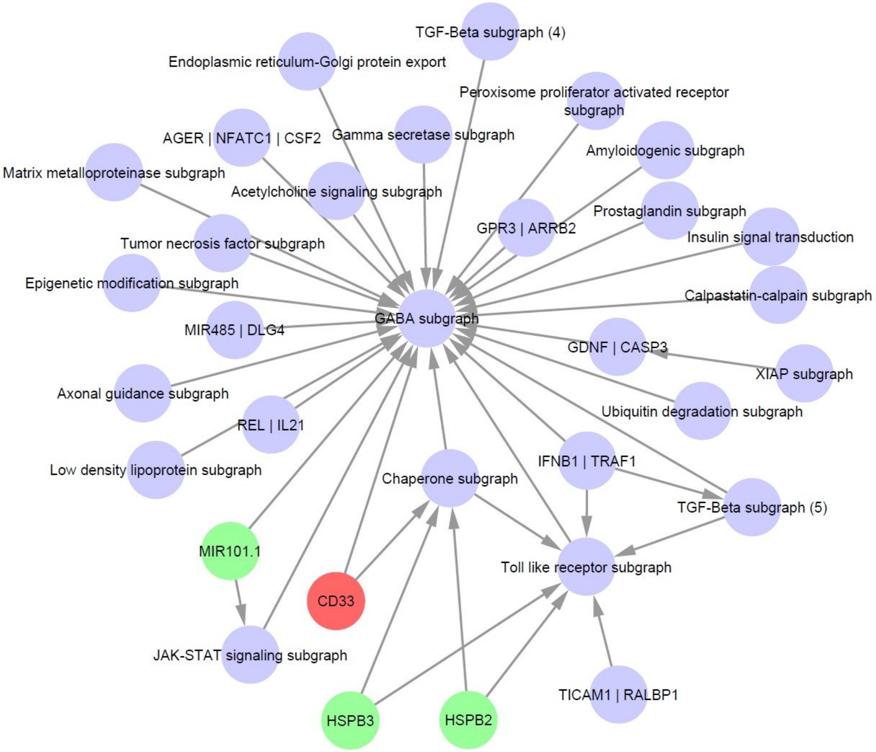


**Supplementary Figure S1**: **Clustered knowledge graph.** Knowledge graph modules (clusters) are annotated with significantly enriched (adjusted $p < 0.05$) NeuroMMSig mechanisms. If the genes in a module do not enrich NeuroMMSig terms significantly, symbols of contained genes are reported. If multiple significant enriched terms could be found, the most significant pathway was used for naming the corresponding node. In case that a module contains a single gene, the gene symbol is reported. CD33 is marked in red, while other single genes are displayed in green, and non-single gene modules in purple.

#### Supplementary Note S2: iVAMBNs Knowledge Integration

The encoded values from the HI-VAEs are then used for training of a Modular Bayesian Network. Part of this training is a structure training, which was performed following three different strategies with five scenarios in total. This gives the opportunity to investigate the level of knowledge integration within the Modular Bayesian Network construction. The three different strategies for structure learning are: 1) completely data driven, 2) knowledge informed, and 3) completely knowledge driven. The second one, knowledge informed, was realized in three different scenarios. The differences between all these options are realized with various black- and white list definitions: while there is no restriction of edges between genes in the completely data driven scenario, the whole network structure is pre-defined in the completely knowledge driven one. More details about the defined blacklists and the definition of the different scenarios can be found in the Methods Section.

Once done for every scenario, the structure learning was evaluated based on the 10-fold cross-validated loss on held out test data shown in Figure **S2**. The lowest average cross-validated loss could be observed in one option of the second scenario, the knowledge informed one, where the original knowledge graph is given as initialization for the hill climber, and the knowledge graph is used as a white list in addition. Therefore, we went ahead with this strategy for all further analyses.

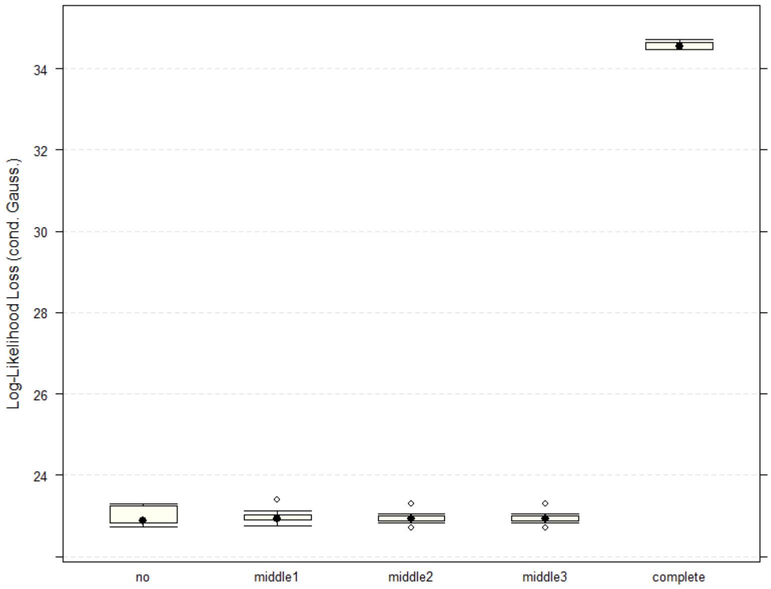


**Supplementary Figure S2**: **Log likelihood loss (i.e. negative log-likelihood) of different knowledge integration strategies.** Completely knowledge driven approach, forcing the BN to have the fixed structure of the knowledge graph, does perform much worse than the completely data driven or knowledge informed approaches. One can observe the lowest average log likelihood loss for the third option of the knowledge informed approach, which forces the KG edges' to be present in the BN, but allows for training of new edges, while using the KG also as the initialization for structure learning.

#### Supplementary Figure S3: Quantitative effect between modules of shortest path.

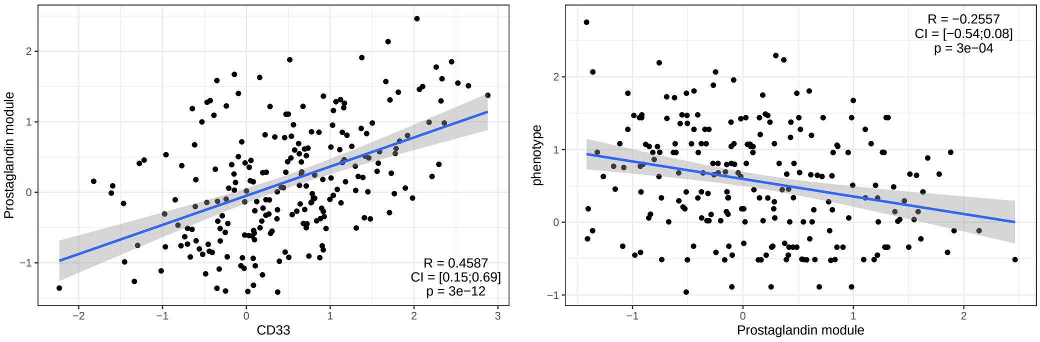


**Supplementary Figure S3: Quantitative effect between modules of shortest path.** Each correlation (R) is shown along with its confidence interval (CI) and multiple testing adjusted p-value. Left: Correlation of CD33 with prostaglandin pathway module. Right: Correlation of prostaglandin pathway module with the phenotype module.

### Supplementary Figure S4: Quantitative effect between modules of newly trained edges with confidence 1

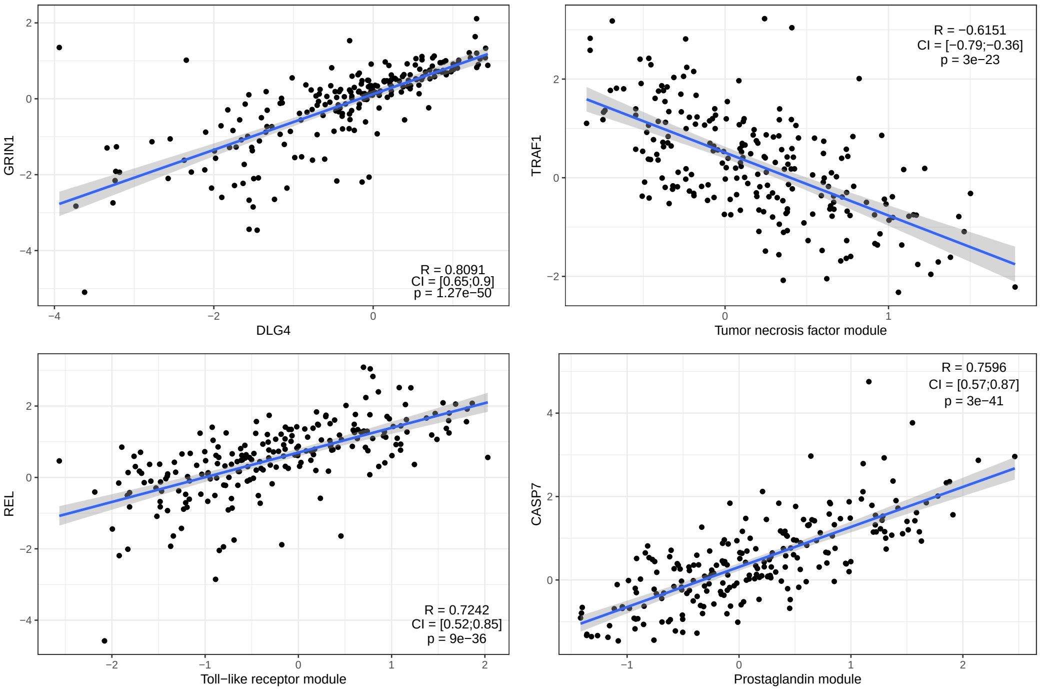


**Supplementary Figure S4**: **Quantitative effect between modules of newly trained edges with confidence 1.** Each correlation (R) is shown along with its confidence interval (CI) and multiple testing adjusted p-value. The *from* module is always shown on x-axis while the *to* module is shown on y-axis.

### Supplementary Table S3: Bootstrap confidence results

A full list of the bootstrap confidence of each possible edge in the Bayesian Network can be found here and in the attached table. For every edge the corresponding start and end note, as well as, the bootstrap strength and the direction is given.

#### Supplementary Figure S5: Overlap of ROSMAP and Mayo network structures

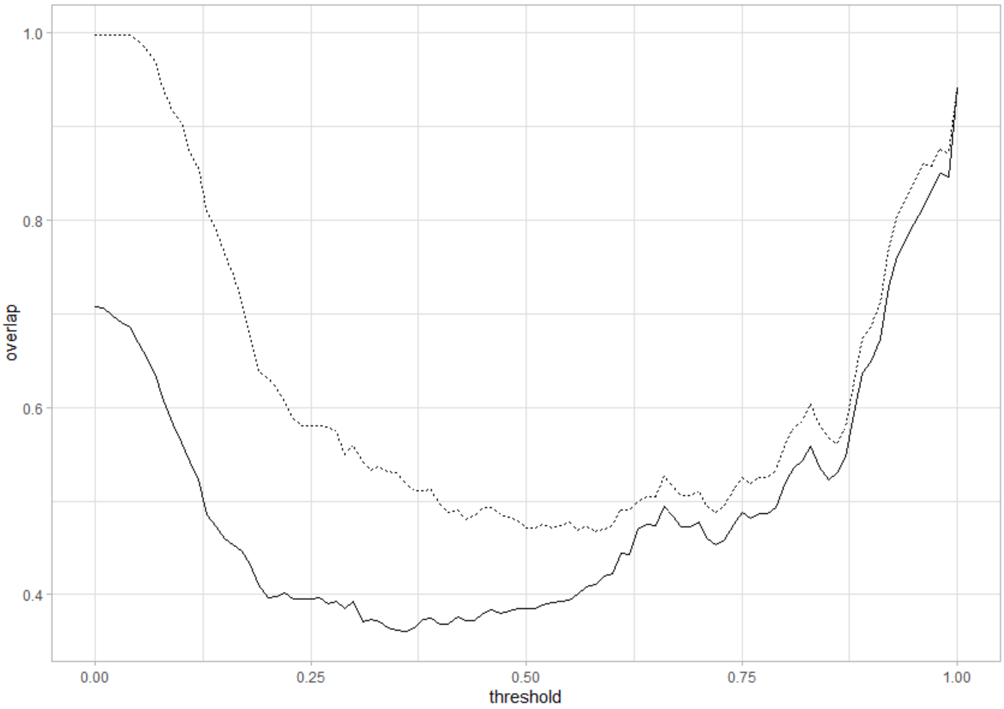


**Supplementary Figure S5**: **Overlap of ROSMAP and Mayo network structures** The overlap of the independent bootstrap structure learning for ROSMAP data and Mayo data is shown for different threshold values. The black line represents the overlap when considering the direction of the edge, the dashed line the overlap of the network skeletons.

#### Supplementary Figure S6: Effects on phenotype scores of up- and down-regulation simulations.

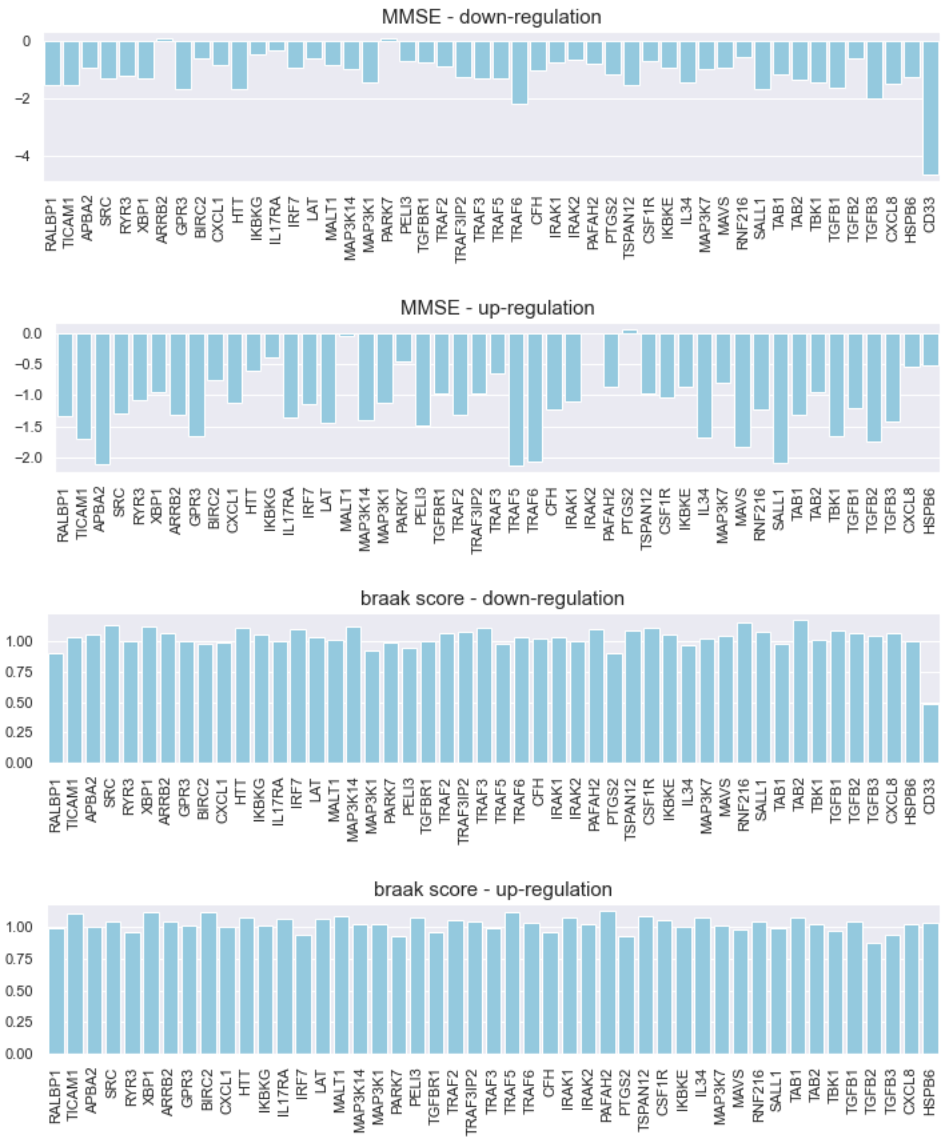


**Supplementary Figure S6**: **Effects on phenotype scores of up- and down-regulation simulations.** The bar plots show the difference between the mean score in the original data and the mean score in the simulated data for each target and each phenotype score, namely MMSE (upper two rows) and Braak score (bottom two rows). First and third row shows the results of under-expression, while second and forth rows shows the results of over-expression.

#### Supplementary Table S4: Graph Clustering Metrics

**Supplementary Table S4**: **Graph Clustering Metrics** The three clustering algorithms: 1) markov clustering, 2) edge betweenness, and 3) infomap were applied on the knowledge graph. For every cluster algorithm the corresponding metrics are given, as well as, the average rank. Printed in bold is the best algorithm according to the respective metric. The algorithms were ranked per metric and the average rank per algorithm was calculated.

| metric | markov | between | infomap |
|---|---|---|---|
| number clusters | **32** | 41 | 40 |
| internal density | 0.1515 | 0.1663 | **0.1662** |
| edges inside | **344.61** | 312.80 | 68.09 |
| av degree | 1.3577 | **1.3708** | 1.0966 |
| expansion | 0.2272 | **0.2141** | 0.4883 |
| cut ratio | **0.0006** | 0.0010 | 0.0015 |
| conductance | **0.0887** | 0.0988 | 0.0987 |
| norm cut | **0.0895** | 0.2078 | 0.2180 |
| average rank | **1.5714** | 1.7143 | 2.7142 |

#### Supplementary Note S3: Evaluating the Model Fit

To evaluate the fit of the overall iVAMBN model we employed the generative nature of our model: Following a topological sorting of the nodes of the DAG of the MBN we first sampled from the distribution of each node conditional on its parent. Notably, for MBN nodes representing modules this amounted to sample from the posterior of the according HI-VAE. Subsequently, the random sample was then decoded via the HI-VAE. We then compared the marginal distribution of each variable based on the synthetic and the real data. Some distributions are shown in Figure **S7**, all other plots are accessible through the github repository.

Furthermore, we compared the correlation matrices of synthetic and real data (cf Figure **S8**). The relative error between the real and simulated values was 0.7138.

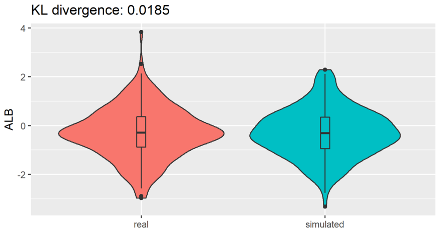


| Type | Mean | SD | 25% | Median | 75% |
|---|---|---|---|---|---|
| *real* | *-0.29* | *1.05* | *-0.89* | *-0.29* | *0.36* |
| *simulated* | *-0.3* | *1* | *-0.95* | *-0.31* | *0.35* |

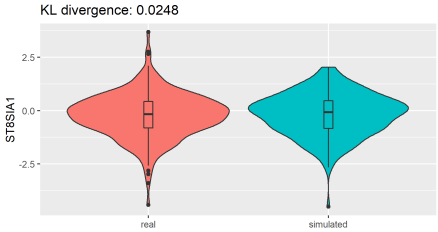


| Type | Mean | SD | 25% | Median | 75% |
|---|---|---|---|---|---|
| *real* | *-0.22* | *1.06* | *-0.81* | *-0.17* | *0.43* |
| *simulated* | *-0.19* | *1* | *-0.83* | *-0.07* | *0.46* |

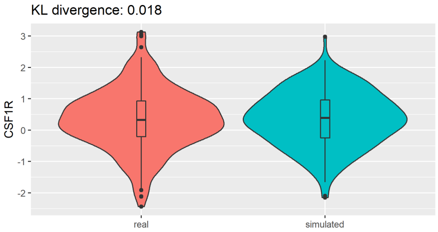


| Type | Mean | SD | 25% | Median | 75% |
|---|---|---|---|---|---|
| *real* | *0.34* | *0.97* | *-0.2* | *0.33* | *0.93* |
| *simulated* | *0.36* | *0.88* | *-0.25* | *0.39* | *0.97* |

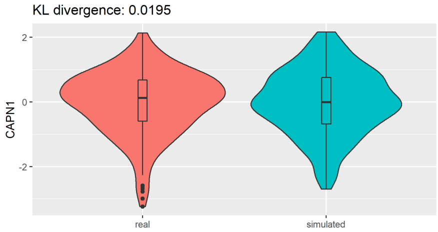


| Type | Mean | SD | 25% | Median | 75% |
|---|---|---|---|---|---|
| *real* | *-0.02* | *0.98* | *-0.59* | *0.13* | *0.68* |
| *simulated* | *-0.01* | *1.01* | *-0.68* | *-0.01* | *0.76* |

**Supplementary Figure S7: Distribution of single feature in real data versus simulated data.**

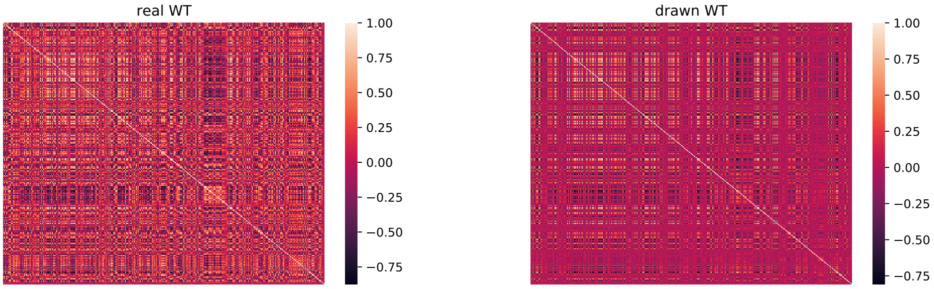


**Supplementary Figure S8: Correlation matrix of real and drawn data.**

#### Supplementary Appendix S1: Github repository of the code for this analysis

The source code used to produce the results and analyses presented in this manuscript are available on a GitHub repository at `https://github.com/traschka/iVAMBN`.
